# Construction of redesigned pMAL expression vector for easy and fast purification of active native antimicrobial peptides

**DOI:** 10.1101/2021.05.26.445771

**Authors:** Lazar Gardijan, Marija Miljkovic, Mina Obradovic, Branka Borovic, Goran Vukotic, Goran Jovanovic, Milan Kojic

## Abstract

Many protein expression and purification systems are commercially available to provide a sufficient amount of pure, soluble and active native protein, such as the pMAL system based on *E. coli* maltose binding protein tag (MBP). Adding specific amino acid tags to the N- or C-terminus of the protein increases solubility and facilitates affinity purification of proteins. However, many of expressed tagged proteins consequently lose functionality, particularly small peptides such as antimicrobial peptides (AMPs). Objective of this study was to redesign the pMAL expression vector in order to increase the efficacy of MBP tag separation from native peptides. Redesign of the pMAL expression vector included introduction of the His_6_ tag and the enterokinase cleavage site downstream from the original MBP tag and Xa cleavage site enabling purification of native and active peptide (P) following two-step affinity chromatography. In the first step the entire MBP-His_6_-P fusion protein is purified through binding to Ni-NTA agarose. In the second step, the purification was performed by adding mixture of amylose and Ni-NTA agarose resins following cleavage of the fusion protein with active His_6_ tagged enterokinase. This removes MBP-His_6_ and His_6_-enterokinase leaving pure native protein in solution. The redesigned pMAL vectors were optimized for cytoplasmic (pMALc5HisEk) and periplasmic (pMALp5HisEk) peptides expression. Two-step purification protocol was successfully applied in purification of active native AMPs, lactococcin A and human β-defensin. Taken together, we established the optimal conditions and pipeline for overexpression and purification of large amount of native peptides, that can be implemented in any laboratory.

## Introduction

Proteins play crucial and diverse roles in all organisms. They represent the building blocks of every cell, act as enzymes, hormones, cell attachment anchors, regulators and source of amino acids, play role in signaling and defense as antibodies or antimicrobial peptides (AMPs), and contribute in many other processes. The protein over-production and purification is a powerful tool in biotechnology and basic science research and need for a variety of purified native proteins is elevated over the last decade [1, 2]. However, the biotechnology industry is still faced with insufficient production capacity due to increasing demand for biologically active proteins and in particular those used as therapeutics [3]. The favorite host for recombinant protein production is an *Escherichia coli*, not only for ability to grow rapidly and achieve high density on cheap media, but also because of well-characterized genetics and the available number of cloning vectors and mutant host strains [4–6]. There are many protein expression and purification systems developed based on *E. coli* host and the corresponding expression vectors. Majority of expression vectors are designed to introduce the amino acids tag to the protein of interest leading to expression of the tagged fusion proteins [6–8]. Many of those fusion proteins retain the activity close to wild type, even in the presence of the tag. However, a significant number of tagged proteins lose the activity and so require to be purified in its native form. This is particularly true for the smaller proteins such as peptides, or more specifically, the antimicrobial peptides (AMPs). Importantly, AMPs are at the forefront of the effort to replace/substitute the antibiotics and tackle the ever increasing antimicrobial resistance problem [9–11].

In this work, we redesigned the pMAL vector which is part of the well known protein expression and purification system typically used to purify insoluble proteins [12–14]. Here, the relatively large size maltose binding protein (MBP) of ~43 kDa is fused as a tag to the protein of interest and the native protein is purified following overexpression and purification through amylose resin column and cleavage by Xa protease which cleavage site is positioned between the MBP and the protein. This system is designed to require the use of the size exclusion columns/filters to separate the many smaller proteins from MBP. However, if the native protein of interest is too small, chemical characteristics of the size exclusion columns/filters, cause non-specific binding of the small peptide to the column. Therefore, the rational to redesign the pMAL vector and system was to accommodate the vector and develop the protocol for overexpression and purification of the small peptides, specifically AMPs. The effectiveness of novel system was tested using well characterized AMPs, lactococcin A (LcnA) and human β-defensin (hBD).

## Materials and Methods

### Bacterial strains and growth conditions

*Escherichia coli* strains DH5α [15] and ER2523 (New England Biolabs, Ltd., UK) were grown in Luria Bertani (LB) medium at 37°C with aeration (180 rpm). *Lactococcus lactis* BGMN1-596 was grown in M17 medium (Merck GmbH, Darmstadt, Germany) supplemented with D-glucose (0.5% w/v) (GM17) at 30°C [16]. Solid medium and soft agar were made by adding 1.5% and 0.7% (w/v) agar (Torlak, Belgrade, Serbia) to the liquid media, respectively. Ampicillin (100 μg/ml) was used for selection and maintaining of transformants. Isopropyl-β-D-1-thiogalactopyranoside (IPTG; Serva, Heidelberg, Germany) in appropriate concentrations was used for the induction of protein expression.

### DNA manipulations

For plasmid isolation from *E. coli* transformants, a Thermo Fisher Scientific GeneJET Plasmid Miniprep kit was used according to the manufacturer’s recommendations (Thermo Scientific, Lithuania). Digestion with restriction enzymes was conducted according to the supplier’s instructions (Thermo Fisher Scientific Waltham, MA, USA). DNA was ligated using T4 DNA ligase (Agilent technologies, USA) according to the manufacturer’s recommendations. Standard heat-shock transformation method was used for transformation of *E. coli* with a plasmid [15].

### Site-directed mutagenesis

Desired mutations were introduced as described previously by Miljkovic *et al*. [17] using oligonucleotide primers (and their reverse complements) carrying mutations (see Table 1). PCR amplification of each plasmid DNA strand for site-directed mutagenesis was performed separately by adding only one primer. Amplification was conducted by Phusion High Fidelity DNA Polymerase (Thermo Fisher Scientific) one min/kb. Single strand amplicons obtained by forward and complement primers were annealed (see below) and digested with 1 μl (10 U) of *Dpn*I restriction enzyme per reaction (at 37°C for 2 h) to destroy the methylated template strands. Non-digested plasmid DNA was used for transformation of *E. coli* DH5α high-competent cells by heat shock treatment. Transformants were selected on LA Petri dishes containing ampicillin. To obtain plasmid DNA from the selected colonies, the GeneJET plasmid miniprep kit was used according to the manufacturer’s recommendations (Thermo Fisher Scientific). The introduction of the desired mutations was confirmed by sequencing (Macrogen Europe, The Netherlands).

**Table 1.**
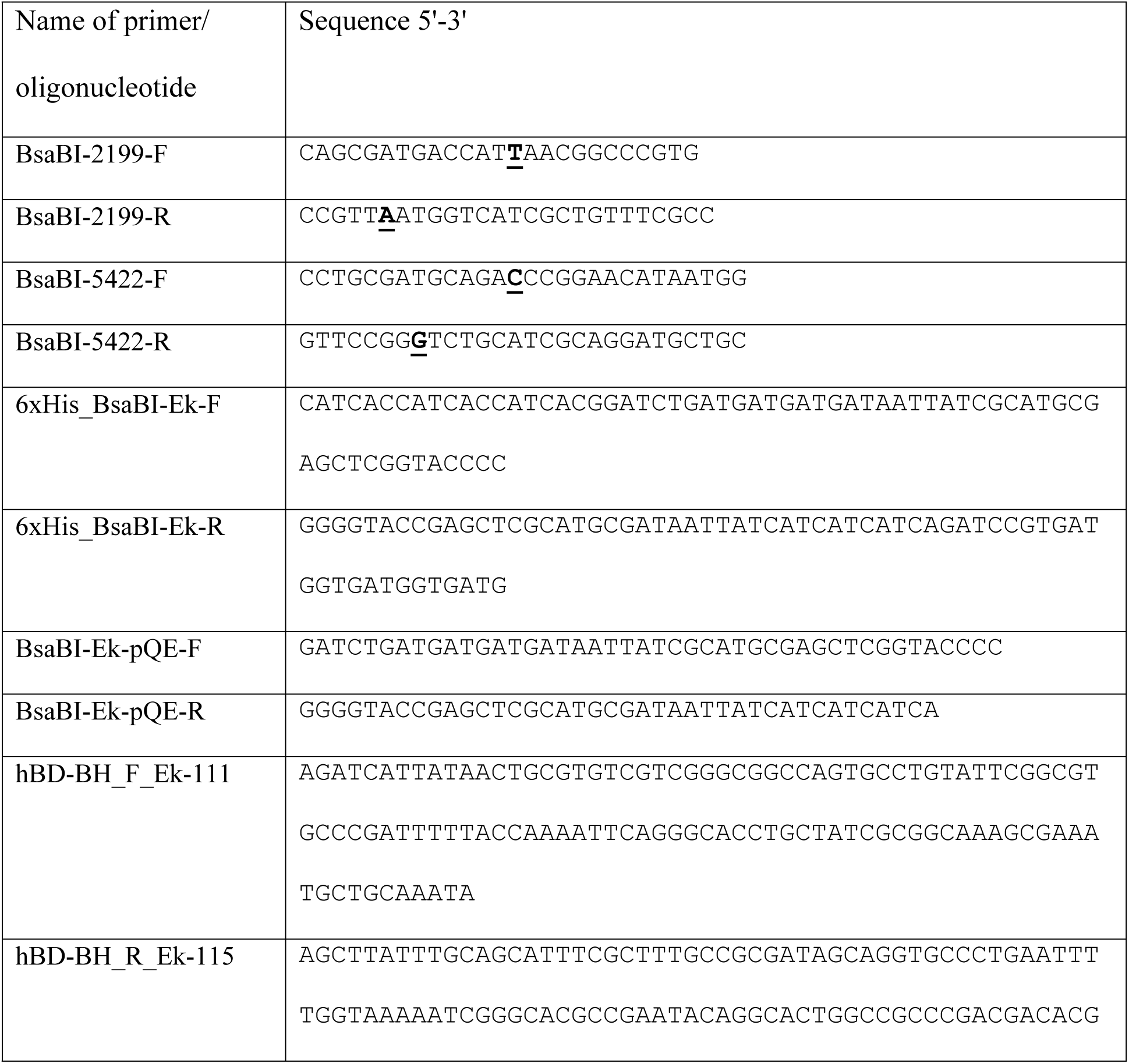
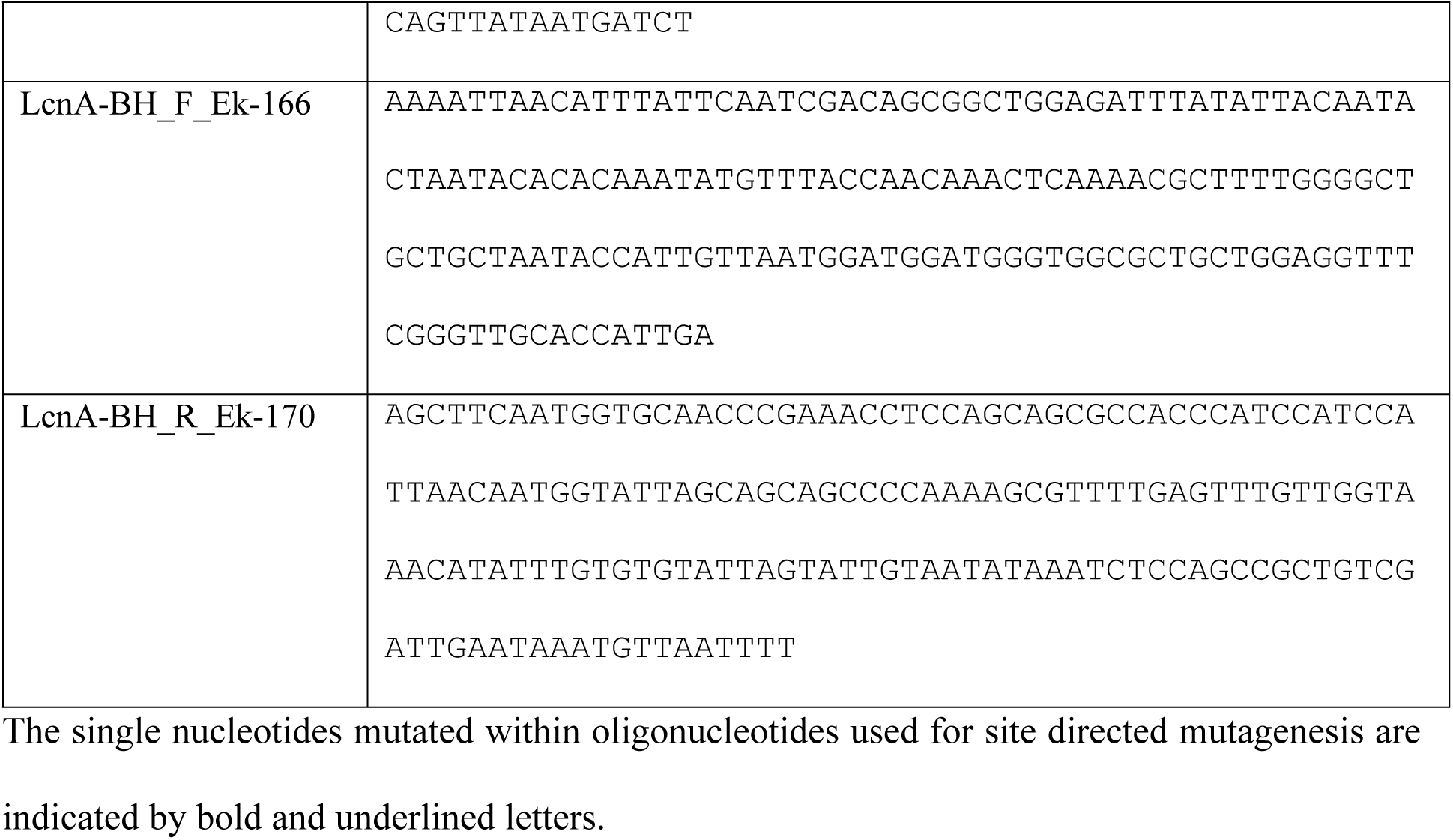
Primers and olignucleotides used in this study.

### Annealing of DNA oligonucleotides

Annealing of two single-stranded DNA oligonucleotides with complementary sequences was done by heating and cooling method. Oligonucleotides were dissolved in pure water. Annealing mixture (50 μl) composed of two oligonucleotides (with the equimolar concentration 10 μM) in annealing buffer (10 mM Tris-HCl-pH 7.5, 50 mM NaCl, 1mM EDTA) was incubated for 5 min at 95-100°C and after that slowly cool down to room temperature before transferring it on ice. Hybridized oligonucleotides were used for ligation with predigested vectors or stored at −20°C.

### Overexpression and two-step purification of the recombinant AMPs in *E. coli* ER2523

In this study we tested the efficacy of newly constructed (reconstructed) pMAL expression vectors pMALc5HisEk and pMALp5HisEk by expressing and purifying two AMPs, through two-step purification. We used Ni-NTA agarose and amylose resins affinity chromatographies in two different combinations. Transformants of *E. coli* ER2523 were maintained overnight on LA Petri dishes containing ampicillin (100 μg/ml) and glucose (1%) at 30°C. The next day new cultures (each 200 ml of LB with 1% glucose) were inoculated using 2% sample of overnight culture and incubated at 30°C with aeration (180 rpm on rotatory shaker). Expression of recombinant peptides was carried out in LB containing ampicillin (100 μg/ml) and glucose (1%) and protein production was induced in logarithmic growth phase (OD_600_= 0.8-1.0) with addition of 0.3 mM (final) IPTG for 3 h. Bacterial cells were collected by centrifugation at 4500 x *g* and before purification the level of induction was tested by comparing the amount of total proteins from the same amount of induced and non-induced cells. Purification including cell lysis, affinity chromatographies, and cleavage of the fusion protein with enterokinase that were performed according to manufacturer’s instructions (amylose resin purification – according to pMAL Protein Fusion & Purification System, New England Biolabs, Ltd., UK; Ni-NTA agarose resin purification – according to The QIAexpressionist, Qiagen Gmbh, Hilden, Germany; cleavage with recombinant bovine enterokinase – according to GenScript, New Jersey, USA), with the addition of cell lysis step that was performed in column buffer (CB) with 1 mg/ml lysozyme for 30 min on ice.

Firstly, the Ni-NTA agarose resin purification step was performed and then purified fusion protein was digested in 1 X enterokinase buffer (20 mM Tris-HCl, pH 7.4, 50 mM NaCl, 2 mM CaCl_2_) using 10 U of Recombinant Bovine His_6_-Enterokinase (GenScript, Piscataway, NJ, USA) overnight at room temperature. In order to place the purified protein in 1 X enterokinase buffer, the buffer exchange was done using Amicon Ultra 0.5 ml 30K Centrifugal filters (Merck Millipore Ltd, Cork, Ireland). After 24 h of digestion, total cleavage of the purified protein was obtained and the second purification step using amylose and Ni-NTA agarose resins mix was performed according to the instructions (The QIAexpressionist).

Proteins from every step of expression, purification and proteolysis were analyzed on 12.5% sodium dodecyl sulfate polyacrylamide gel electrophoresis (SDS-PAGE). Samples for SDS-PAGE were mixed with 2 X sample loading buffer (125 mM TrisHCl pH6.8, 10 mM EDTA, 4% SDS, 25% glycerol, 5% β-mercaptoethanol, 0.07% bromophenol blue) at 1:1 ratio. Samples were denatured by heating for 5 min at 100°C before loading on the gel.

### Antimicrobial activity assay

A spot-on-the-lawn inhibition assay was used for testing antimicrobial activity of purified recombinant AMPs as previously described [18]. *L. lactis* BGMN1-596 [16] was used as sensitive strain in antimicrobial assay. 5 μl of the purified AMP was applied to soft agar inoculated with *L. lactis* BGMN-596. The presence of inhibition zones was examined after 24 h following incubation at 30°C. A clear zone of inhibition was taken as evidence of antimicrobial peptide activity. The antimicrobial activity assay was performed in at least two independent experiments. The enterokinase buffer (in which the purified AMPs were resuspended) was taken as a negative control in an antimicrobial assay.

## Results and Discussion

### Rational for redesigning pMAL expression vector

In our previous work the most successful expression and purification of AMPs, was achieved with the pMAL expression and purification system [19–21]. The pMAL expression vector (New England Biolabs) provided a high yield of soluble native peptide following precise excision of MBP tag using Xa protease. However, we encountered the problem in separating the MBP tag from dissolved purified low molecular weight AMPs. We tried to facilitate the process by passing digested eluates through a 10 kDa cut-off column, but the yield was not sufficient due to the high binding of the small peptide to the column, even though the low-binding columns were used.

To meet the requirements for the fast and simple separation of expressed AMPs from the MBP tag we decided to design and generate an improved version of pMAL expression vector, but different from the one already available pDEST-HisMBP (www.addgene.org, plasmids #11085). pDEST-HisMBP contains the His_6_ tag positioned at the MBP N-terminus and a TEV protease recognition site at C-terminus for the excision of the recombinant peptide [22]. There is a commercially available His_6_ tagged enterokinase protease that cuts immediately after the recognition cleavage site potentially releasing precisely the expressed native peptide. The idea was to use pMAL vector and introduce the His_6_ tag and the enterokinase cleavage site downstream from the MBP and Xa cleavage site and just upstream of the Multi Cloning Site (MCS) used for cloning the desired peptide DNA (Fig 1). This will provide the effective affinity separation of the MBP-His_6_ tag and the His_6_-enterokinase from the peptide following cleavage of the composite MBP-His_6_-Ek-P fusion protein with the His_6_-enterokinase and use of Ni-NTA agarose.

**Fig 1.**
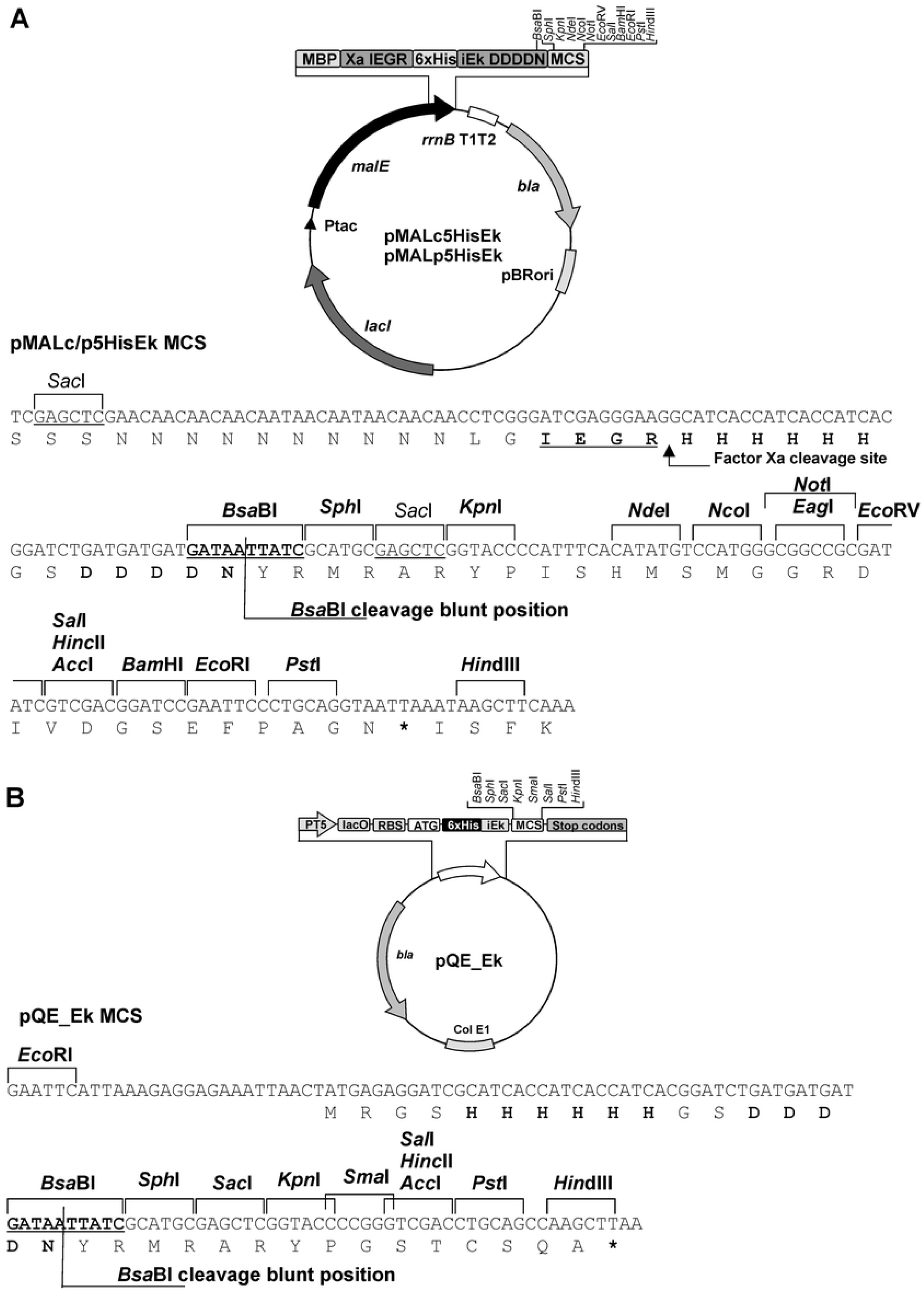
Construction of redesigned pMAL and pQE vectors. Circular maps and depicted redesigned nucleotide sequence of (A) pMALc/p5HisEk and (B) pQE_Ek vectors. Arrows indicate the size and direction of gene transcription. Unique restriction sites are indicated by bold letters. *malE:* maltose binding protein (MBP) gene, Xa: Xa factor cleavage site IEGR (typical Xa cleavage site is IE/DGR), 6xHis: 6xHis tag sequence, iEk: incomplete enterokinase (Ek) cleavage site (DDDDN) to be reconstituted into complete Ek cleavage site DDDDK following cloning, MCS: multiple cloning site with restriction sites indicated, *bla:* ampicillin resistance gene, *lacI, lacI*^q^ repressor gene, *Ptac: taq* promoter, pBRori: pBR322 origin of replication, PT5: T5 promoter, lacO: *lac* operator, RBS: ribosome-binding site, ATG: start codon, Stop Codons: stop codons in all three reading frames, Col E1: ColE1 origin of replication.

### Construction of redesigned pMAL and pQE expression vectors

We constructed redesigned pMAL expression vectors pMALc5HisEk and pMALp5HisEk based on pMALc5X (for expression of proteins in the cytoplasm; New England Biolabs) and pMALp5X (for expression of proteins in the periplasm; New England Biolabs). Namely, the MBP at the N-terminus in pMALp5X contains the leader peptide for transfer of synthesized protein to the periplasm. The restriction enzyme recognition site sequence analysis showed that the best restriction enzyme DNA sequence that can form the enterokinase recognition site (DDDDK) and be placed upstream of the 5’ end from the MCS is *BsaB*I (GATNN/NNATC) (see Fig 1). The choice of *Bsa*BI restriction enzyme was further supported by the fact that it is a very efficient restriction enzyme that cuts DNA at 60-65°C, with no star activity, introducing blunt ends. However, the sequence analysis of pMALc5X and pMALp5X vectors revealed that they possess two *Bsa*BI restriction sites; one in the *malE* gene (position 2199c/2273p, respectively) and the other at position 5422c/5497p, respectively). To destroy these sites so that the *Bsa*BI restriction site could be introduced upstream of the multi cloning site for cloning DNA fragments, the site-specific mutagenesis of both *Bsa*BI restriction sites in pMAL vectors was done using the primer pairs listed in Table 1. Since *Bsa*BI restriction sites are mapped in coding genes, vital for plasmid maintenance and expression, the mutations introduced by site directed mutagenesis (the codons showing the highest frequency in *E. coli* were selected) formed the same sense codons. The success of the restriction site mutagenesis was first checked by digestion with *Bsa*BI restriction enzyme, and finally confirmed by sequencing. Before proceeding to further construction, the functionality of the mutated pMAL vectors (pMALc5XΔBsaBI and pMALp5XΔBsaBI) was checked. We induced the MBP expression using IPTG and detected the production of overexpressed MBP protein with the correct molecular weight on PAGE SDS electrophoresis.

The His_6_ tag and the *Bsa*BI restriction site were introduced upstream of the multi cloning site of either pMALc5XΔBsaBI or pMALp5XΔBsaBI using oligonucleotides 6xHis_BsaBI-Ek-F and 6xHis_BsaBI-Ek-R (see Table 1). After annealing, double strand hybrid DNA fragment was inserted into the *Xmn*I restriction site. In addition to *Bsa*BI site, oligonucleotides were designed to introduce an extra *Sac*I restriction enzyme recognition site downstream from *Bsa*BI (Fig 1A). The *Sac*I restriction enzyme digestion was used to check for the correct orientation of the inserted DNA fragment. The plasmid DNA from transformants was isolated and digested with the *Sac*I restriction enzyme. Clones that produced a fragment of 104 bps were selected because they contain a cloned fragment in the right orientation. This was confirmed by sequencing. We intentionally introduced the incomplete enterokinase recognition site (iEk: DDDDN instead of complete one DDDDK) being able to manipulate the N nucleotide in *Bsa*BI restriction site [introducing asparagine (N) instead of Lysine (K)] (see Fig 1A). The reason for this is to emphasize need to add an extra adenine to the 5’ end of the desired protein/peptide coding sequence (or to the forward primer used for amplification of desired protein/peptide coding sequence) in order to obtain both, in frame protein/peptide synthesis with MBP tag and the constitution of complete enterokinase cleavage site (DDDDK). Two newly constructed expression vectors were named pMALc5HisEk and pMALp5HisEk. The inducibility and production of overexpressed MBP-His_6_ from the redesigned pMAL vectors was checked following induction with 0.3 mM IPTG for 3 hours, and so the production of correct fusion MBP-His_6_ protein was obtained (Fig 2A).

**Fig 2.**
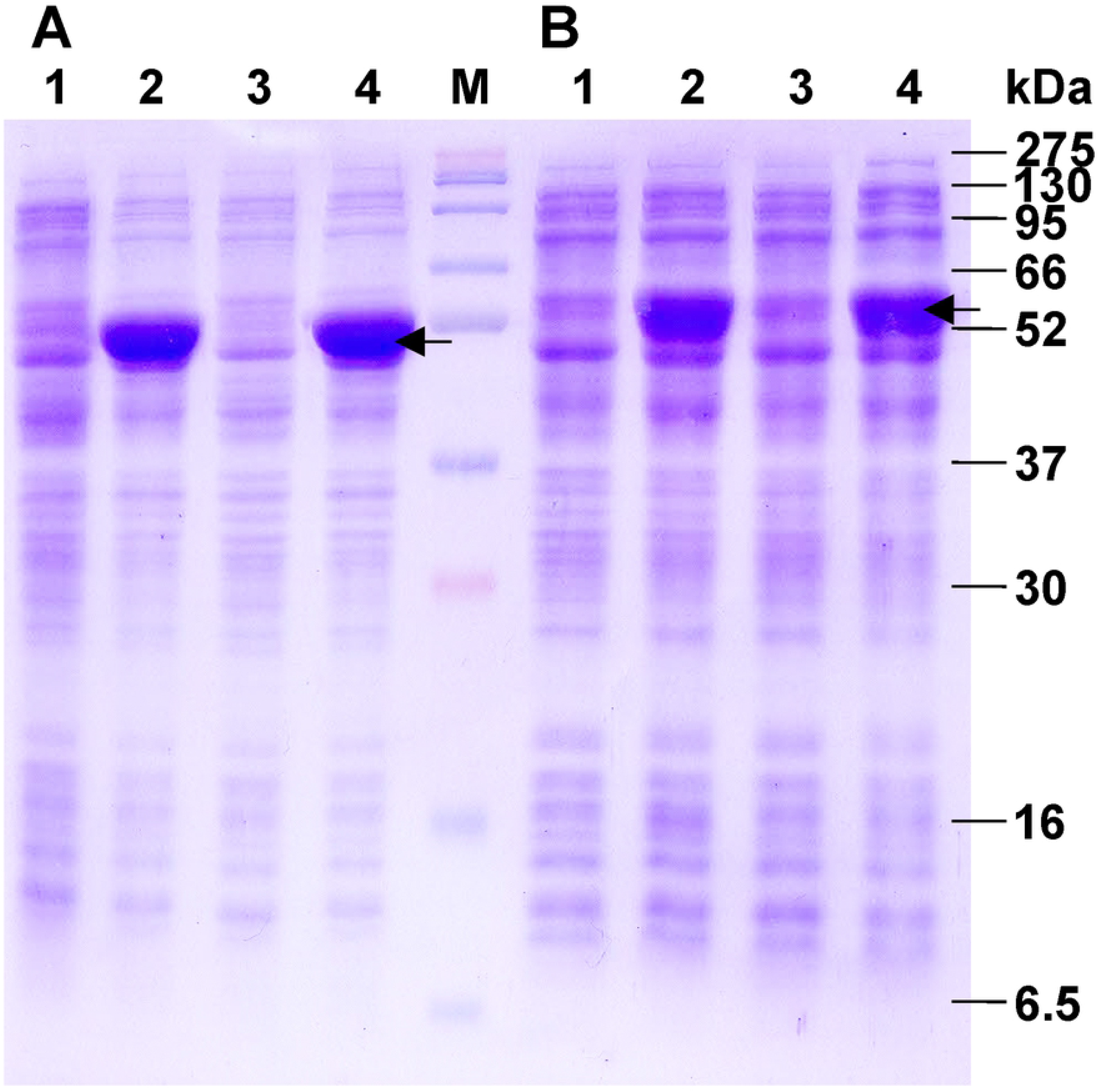
Expression quality control from pMALc5HisEk and pMALp5HisEk vectors. SDS-PAGE analysis of (A) MBP-His_6_ expression in *E. coli* DH5α and (B) recombinant fusion proteins MBP-His_6_-LcnA and MBP-His_6_-hBD expression in *E. coli* ER2523, before and after induction. (A) Total proteins of *E. coli* DH5α carrying pMALc5HisEk or pMALp5HisEk before (lane 1 and 3) and after induction with 0.3 mM IPTG (lane 2 and 4). Note a significant increase of MBP-His_6_ cytoplasmic (lane 2) and periplasmic (lane 4) double tag fusion protein expression (~43 kDa; indicated by black arrow). (B) Total proteins of *E. coli* ER2523 carrying pMALc5HisEk_LcnA1 or pMALp5HisEk_hBD1 before (lane 1 and 3) and after induction with 0.3 mM IPTG (lane 2 and 4). Note a significant increase in expression of cytoplasmic fusion protein MBP-His_6_-LcnA (lane 2; ~50 kDa;) and periplasmic fusion protein MBP-His_6_-hBD (lane 4, ~50 kDa), indicated by black arrow. Lane M: protein molecular weight marker (BlueEasy Prestained Protein Marker)

In addition, we constructed pQE_Ek expression vector for the N-terminal peptide fusions following the same strategy and using BsaBI-Ek-pQE-F and BsaBI-Ek-pQE-R oligonucleotides (see Table 1) and inserting a DNA fragment carrying the incomplete enterokinase cleavage site including *Bsa*BI restriction site into pQE30 previously digested with *Bam*HI (Fig 1B).

### Optimization of expression and purification of LcnA and hBD AMPs using redesigned pMAL vectors

To demonstrate the efficacy of redesigned pMAL vectors pMALc5HisEk and pMALp5HisEk, two AMPs were selected for overexpression, purification and functionality check: lactococcin A (LcnA) for cytoplasmic expression and human beta defensin (hBD) for periplasmic expression, to allow a formation of disulfide bridges.

The coding sequences for LcnA and hBD were synthetic and obtained by annealing of corresponding oligonucleotides (Table 1). The codon selection for hBD was optimized for codon usage of *E. coli.* Extra adenine (A) were added upstream of coding sequences for both AMPs while 3’ ends contained the sticky ends compatible with *Hind*III in digested vectors (Table 1). The dsDNA fragments obtained by annealing of corresponding oligonucleotides were cloned into *Bsa*BI-*Hind*III pre-digested pMALc5HisEk or pMALp5HisEk vectors. More than 90% of analyzed transformants contained plasmid carrying the desired DNA fragment. We sequenced 5 clones from each transformation using pMalE forward primer to confirm the orientation, in frame position with the MBP-His_6_ tags, and the presence of the correct enterokinase cleavage recognition site. The newly generated pMALc5HisEk_LcnA1 and pMALp5HisEk_hBD1 constructs were stored at −80°C in LB containing 15% glycerol and further used for the expression and purification experiments.

For overexpression of LcnA and hBD AMPs, the *E. coli* strain ER2523 was transformed with pMALc5HisEk_LcnA1 or pMALp5HisEk_hBD1 and grown overnight at 30°C. Next day the production of AMPs was induced in the logarithmic phase of growth and the level of induction was tested comparing the amount of total proteins from the same amount of induced and non-induced cells (Fig 2B).

Next, we demonstrated that both tags are individually efficient/active, enabling purification of overexpressed MBP-His_6_-LcnA and MBP-His_6_-hBD. Induced cultures were divided into two equal aliquots and expressed fusion proteins were purified either via amylose resin (binding MBP tag), or via Ni-NTA agarose resin (binding His_6_). The amount and purity of MBP-His_6_-LcnA and MBP-His_6_-hBD purified via MBP or His_6_ tag were comparable (S1 and S2 Fig).

In the first strategy of two-step purification, first step was use of amylose resin to pull down MBP-His_6_-AMP fusion protein, followed by cleavage with enterokinase, and incubation with Ni-NTA agarose resin in the second step to remove the MBP-His_6_ and His_6_-enterokinase from solution, leaving native AMP purified. However, there was no binding of the MBP-His_6_ to the Ni-NTA agarose resin (Fig 3) in three independent attempts. This implied that part of the MBP-His_6_ fusion protein is most likely is not available for immobilization on Ni-NTA agarose due to irreversible change in MBP conformation after maltose binding [23]. Therefore, we changed the order of affinity matrixes application to solve this problem.

**Fig 3.**
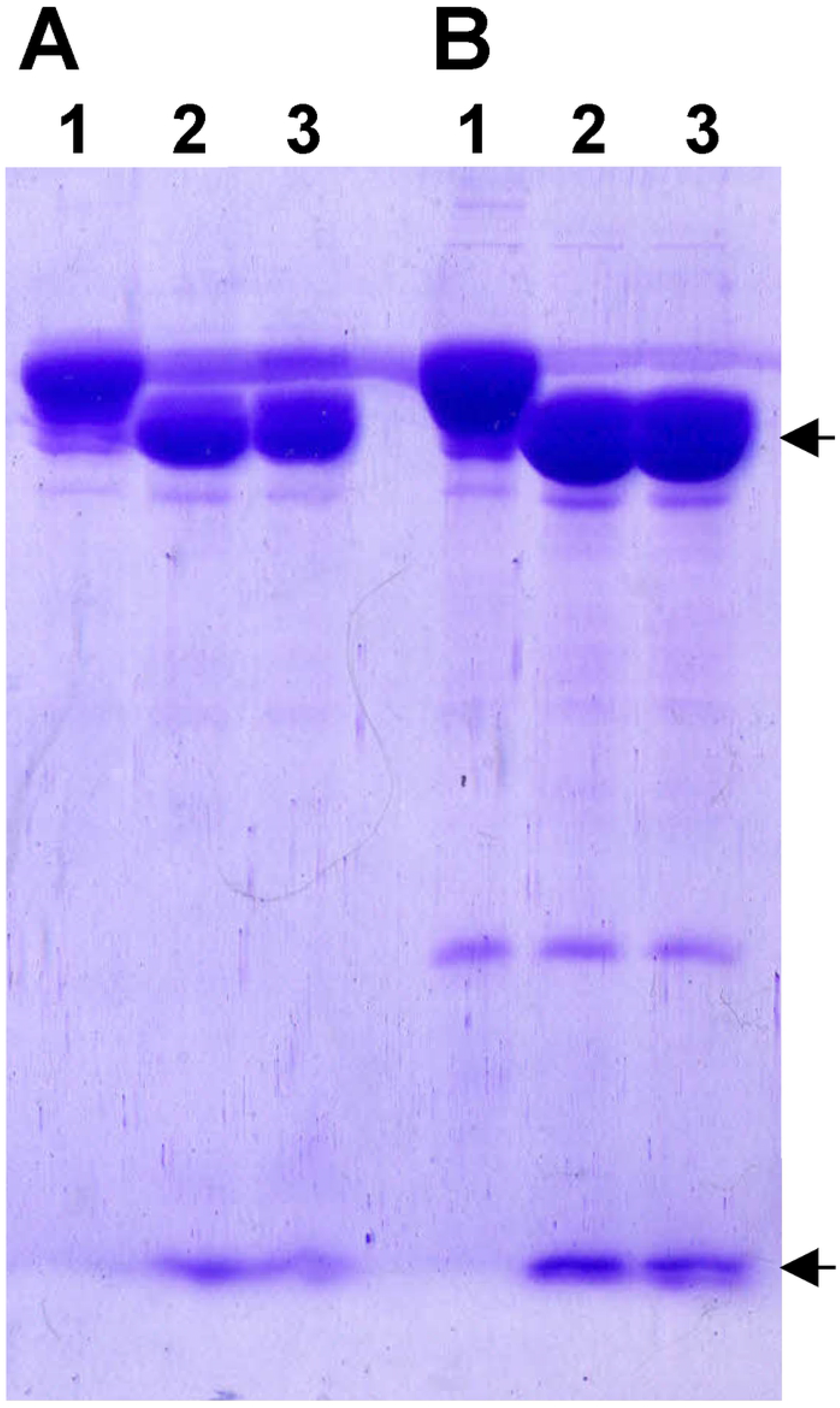
Failing attempt of LcnA and hBD AMPs purification using subsequent application of amylose and Ni-NTA agarose resins. SDS-PAGE analysis of two-step protein purification of (A) LcnA and (B) hBD AMPs, using first amylose and then Ni-NTA agarose resins. (A) Lane 1: eluted fusion protein MBP-His_6_-LcnA using amylose resin, (B) Lane 1: eluted fusion protein MBP-His_6_-hBD using amylose resin, (A and B) Lane 2: products obtained after fusion protein cleavage with His_6_-enterokinase (enterokinase splits the fusion protein into MBP-His_6_ of ~43 kDa and AMP of ~5 kDa), Lane 3: products obtained after second step Ni-NTA agarose affinity chromatography. Black arrows indicate the products following the cleavage by His_6_-enterokinase and the inability of MBP-His_6_ to bind Ni-NTA agarose (see lane 3) following the second step purification.

In the second strategy/attempt, the splitting of MBP-His_6_ and AMP peptide was performed by overnight digestion of AMP with His_6_-enterokinase at room temperature following purification of using either MBP or His tag, (see Materials and Methods). We noted that the efficiency of the protease cleavage was much higher when protein was isolated using Ni-NTA agarose (S3 Fig, line 3), although the proteins were completely digested after 24 h, differences were observed at earlier time points. This might be the consequence related to difference in the availability of the enterokinase cleavage site present on the original MBP-His_6_-Ek-P fusion protein that has undergone conformational changes due to maltose binding. The removal of MBP-His_6_ and His_6_-enterokinase was accomplished in second step purification using a mix of amylose (to bind MBP-His_6_ tag) and Ni-NTA agarose (to bind His_6_-enterokinase) in a ratio of 10:1 (Fig 4) only from digestion of a fusion protein purified using Ni-NTA agarose in first step. The expressed AMPs, LcnA (Fig 4A, line 5) and hBD (Fig 4B, line 5) were isolated in over 90% purity as shown when compared to the sample obtained after cleavage with enterokinase (Fig 4A and 4B, lane 4).

**Fig 4.**
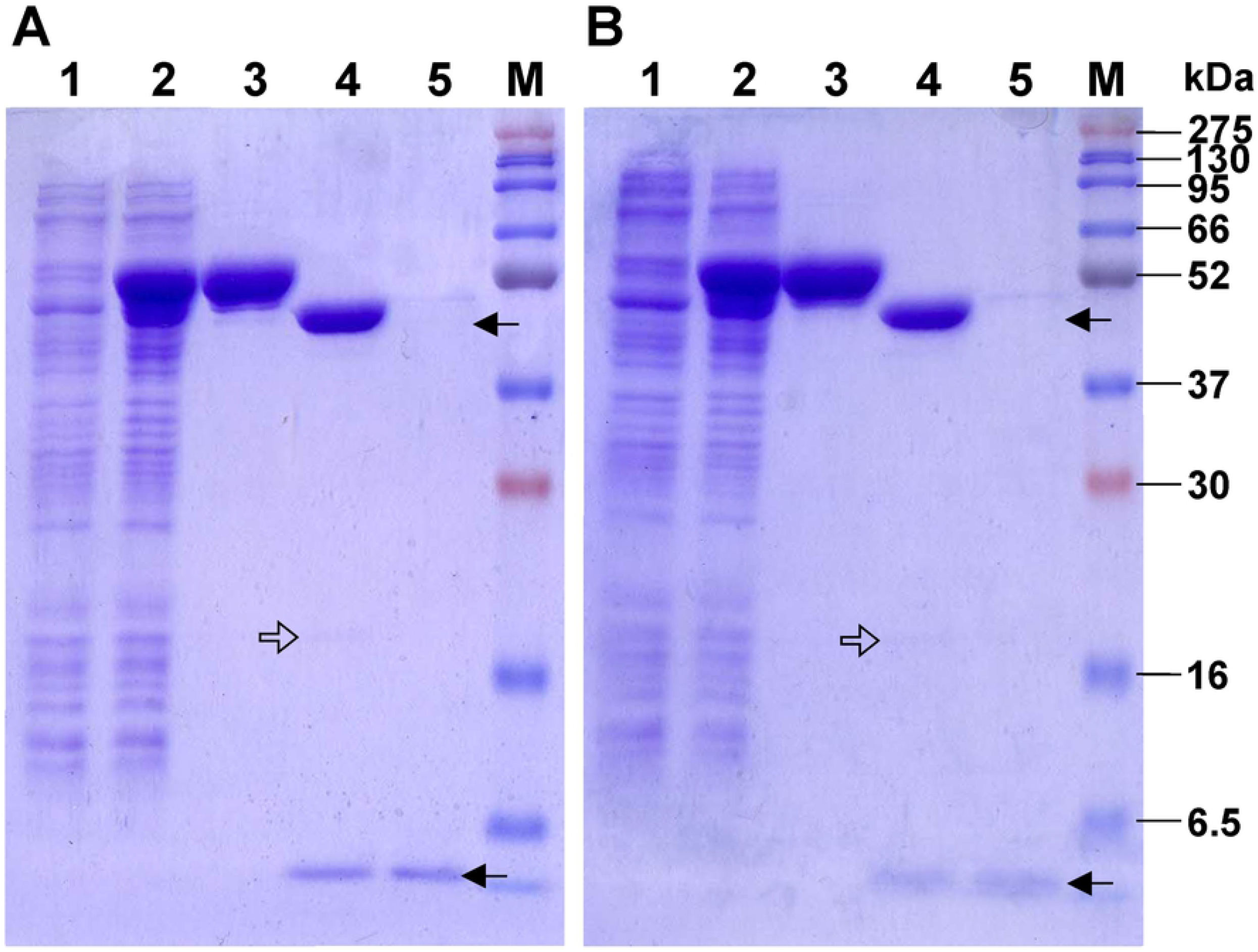
Two-step purification of native LcnA and hBD AMPs using subsequent application of Ni-NTA agarose and amylose : Ni-NTA agarose (10:1) resins mix. SDS-PAGE analysis of two-step protein purification of (A) LcnA and (B) hBD AMPs using first Ni-NTA agarose and then 10:1 mix of amylose and Ni-NTA agarose resins. Lane 1: total proteins of *E. coli* ER2523 carrying (A) pMALc5HisEk_LcnA1 and (B) pMALp5HisEk_hBD1 before induction, Lane 2: after induction with 0.3 mM IPTG (A and B) and expression of ~50 kDa fusion protein, Lane 3: eluted fusion protein (A) MBP-His_6_-LcnA and (B) MBP-His_6_-hBD after purification with Ni-NTA agarose resin, Lane 4: products obtained after cleavage with His_6_-enterokinase which splits fusion protein into MBP-His_6_ tag of ~43 kDa and expressed ~5 kDa AMP (A) LcnA and (B) hBD, Lane 5: purified native AMP without MBP-His_6_ and His_6_-enterokinase after the second step of affinity chromatography purification using 10:1 mix of amylose : Ni-NTA agarose resins. Lane M: protein molecular weight marker (BlueEasy Prestained Protein Marker). Black arrows indicate the products of cleavage by His_6_-enterokinase. Open arrow indicates the position of the His_6_-entrokinase on the gel.

The functionality/bioactivity of the purified native LcnA and hBD was analysed by spot on the lawn antimicrobial assay using *L. lactis* BGMN-596 sensitive strain. Both purified native AMPs showed an antimicrobial activity (Fig 5). The activity of LcnA was higher than hBD and this might be the consequence of incomplete folding of the hBD within hybrid protein in the periplasm or used indicator strain is less sensitive to hBD.

**Fig 5.**
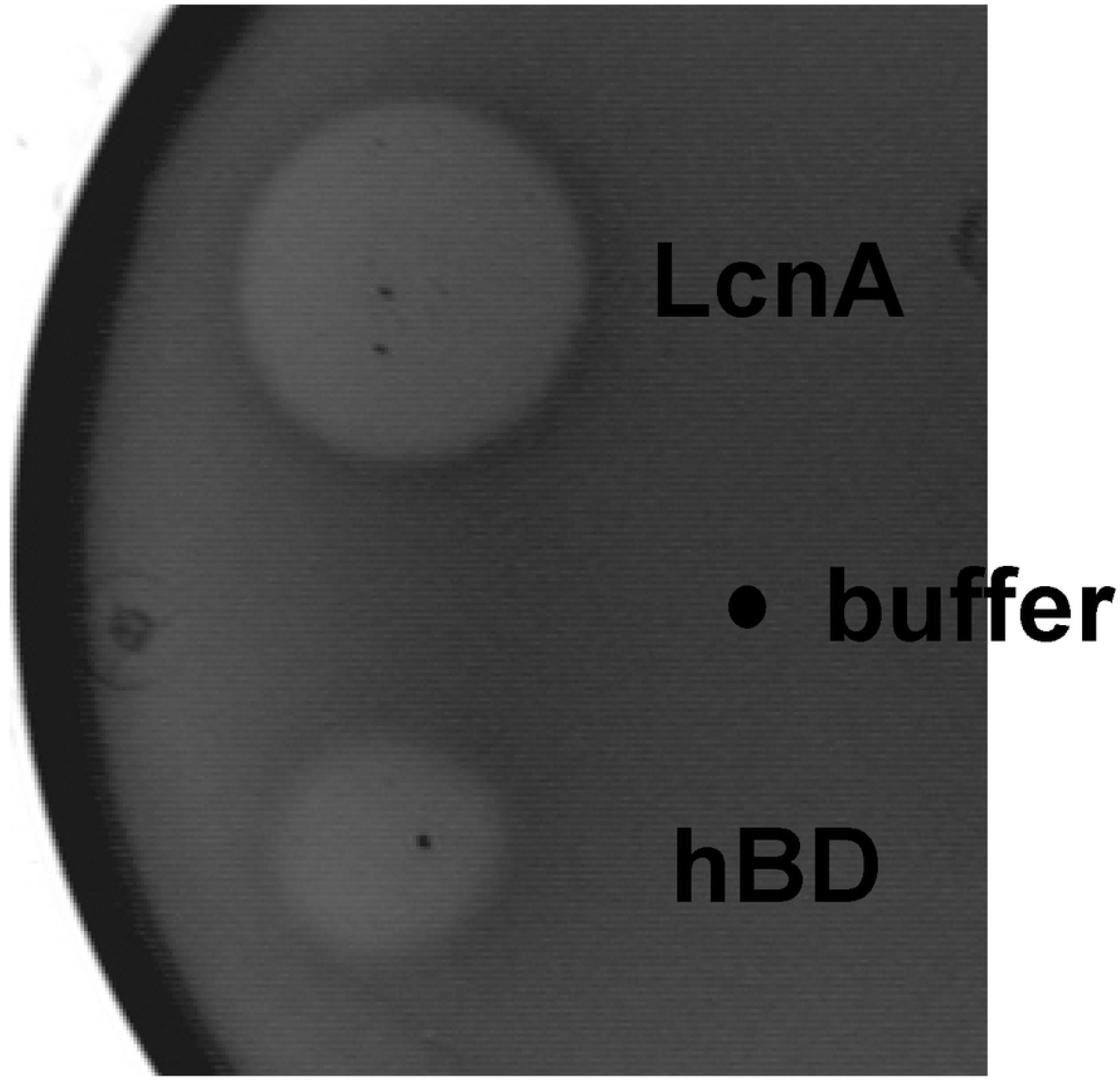
The antimicrobial activity of purified LcnA and hBD. LcnA and hBD were expressed in *E. coli* ER2523 (using pMAL-c/p5HisEk vectors), purified and spotted (5 μl) on *L. lactis* BGMN1-596 sensitive strain. LcnA; lactococcin A, hBD: human beta defensin, buffer: 5 μl of enterokinase buffer (negative control).

## Conclusions

Here we report an efficient procedure for separation of native peptides from the tag(s), using redesigned pMAL expression vector and the two-step affinity chromatography. The idea of redesigning pMAL vector and creating a novel two-tag expression vector(s) was born from the need for rapid purification of larger amounts of active AMPs. We constructed novel pMAL-based vectors, pMALc5HisEk and pMALp5HisEk, which enable purification of active native AMPs from the cytoplasm and periplasm, and demonstrated validity of the novel protocol using LcnA and hBD AMPs. This system is intended for the cloning, expression and purification of heterologous proteins/peptides in *E. coli* and includes fast and reliable procedure for the isolation of active and native recombinant proteins and peptides (Fig 6). By applying this expression and purification protocol, protein purity is greater than 90%, thanks to a double tag separation and protease removal. Moreover, this system is simple and cost-effective, and can be implemented in any laboratory.

**Fig 6.**
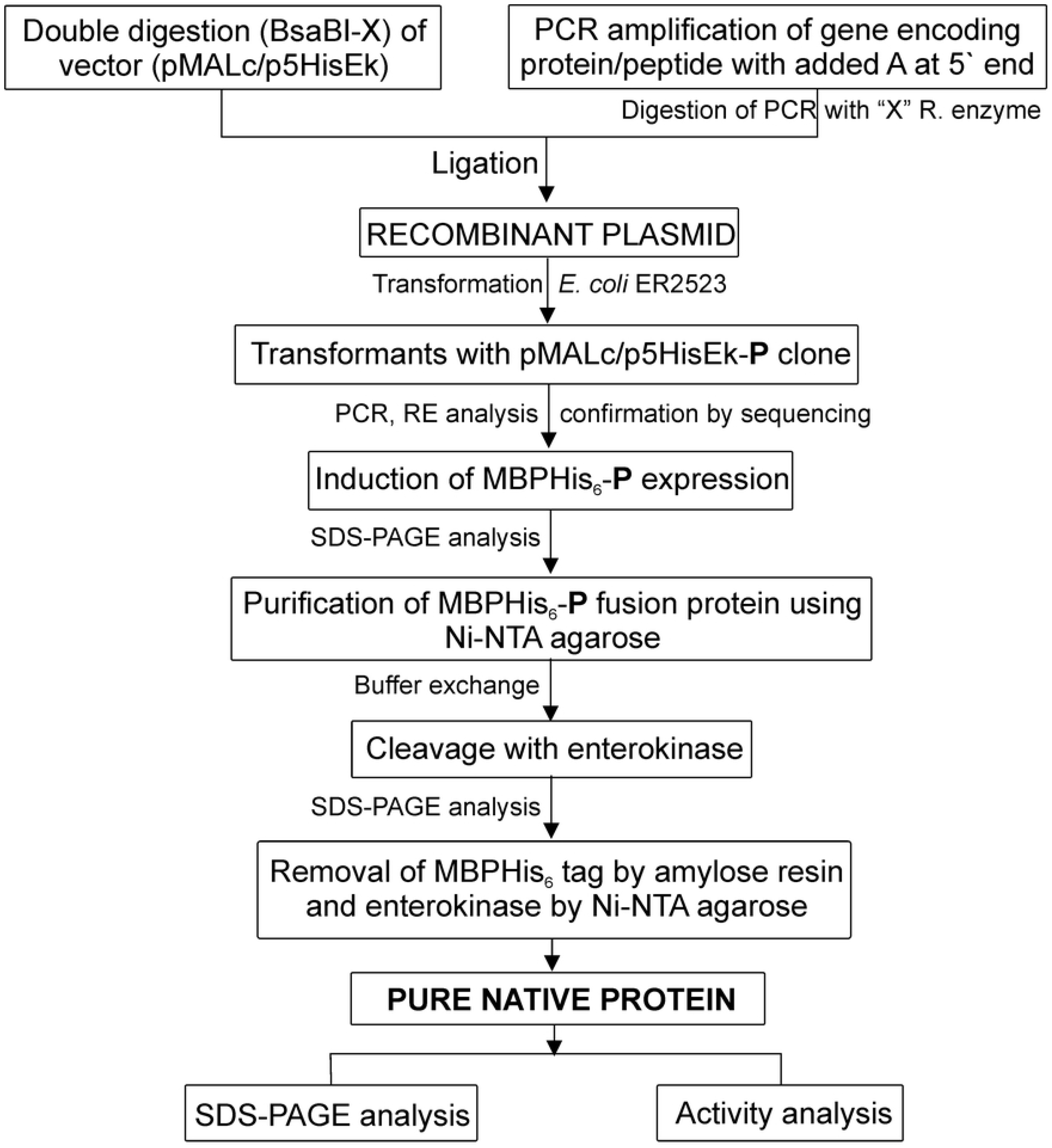
Flow chart presenting pipeline for cloning, expression and two-step purification of peptides/AMPs using redesigned pMAL-c/p5HisEk vectors in *E. coli* ER2523. X: any unique restriction enzyme from the multi cloning site absent from the peptide gene.

## Acknowledgements and Funding

This work was supported by The Ministry of Education, Science and Technological Development of the Republic of Serbia, Republic of Serbia (Grant No.173019).

## Supporting information

**S1 Fig. SDS-PAGE analysis of MBP-His_6_-LcnA recombinant fusion protein purified via (A) Ni-NTA agarose or (B) amylose resin**. Lane 1: flow-through proteins, Lane 2: proteins removed by wash 1, Lane 3: proteins removed by wash 2, Lane 4: proteins removed by wash 3, Lane 5: eluate1, Lane 6: eluate 2, Lane 7: eluate 3, Lane M: protein molecular weight marker (BlueEasy Prestained Protein Marker). Black arrow indicates eluted overexpressed MBP-His_6_-LcnA fusion protein.

**S2 Fig. SDS-PAGE analysis of MBP-His_6_-hBD recombinant fusion protein purified via (A) Ni-NTA agarose and (B) amylose resin.** Lane 1: flow-through proteins, Lane 2: proteins removed by wash 1, Lane 3: proteins removed by wash 2, Lane 4: proteins removed by wash 3, Lane 5: eluate 1, Lane 6: eluate 2, Lane 7: eluate 3, Lane M: protein molecular weight marker (BlueEasy Prestained Protein Marker). Black arrow indicates eluted overexpressed MBP-His_6_-hBD fusion AMP.

**S3 Fig. The cleavage efficacy of enterokinase depends on MBP-His_6_-LcnA purification method.** SDS-PAGE analysis of MBP-His_6_-LcnA recombinant fusion protein after cleavage by His_6_-enterokinase. The MBP-His_6_-LcnA fusion protein purified using Ni-NTA agarose resin (Lane 1) or amylose resin (Lane 2) and cleaved by His tagged entrokinase for 3 h. Lane M: protein molecular weight marker (BlueEasy Prestained Protein Marker). Black arrows indicate the products of enterokinase digestion. Open framed arrow indicates the position of undigested fusion tags-AMP.

## References

1. Vecchio I, Tornali C, Bragazzi NL, Martini M. The Discovery of Insulin: An Important Milestone in the History of Medicine. Front Endocrinol (Lausanne). 2018;9:613. doi: 10.3389/fendo.2018.00613. PMID: 30405529; PMCID: PMC6205949.

2. Geddes BA, Mendoza-Suárez MA, Poole PS. A Bacterial Expression Vector Archive (BEVA) for Flexible Modular Assembly of Golden Gate-Compatible Vectors. Front Microbiol. 2019;9:3345. doi: 10.3389/fmicb.2018.03345. PMID: 30692983; PMCID: PMC6339899.

3. Mergulhao FJM, Monteiro GA, Cabral JMS, Taipa MA. Design of bacterial vector systems for the production of recombinant proteins in *Escherichia coli*. J Microbiol Biotechnol. 2004; 14(1), 1–14.

4. Structural Genomics Consortium; China Structural Genomics Consortium; Northeast Structural Genomics Consortium, Gräslund S, Nordlund P, Weigelt J, Hallberg BM, Bray J, Gileadi O, Knapp S, Oppermann U, Arrowsmith C, Hui R, Ming J, dhe-Paganon S, Park HW, Savchenko A, Yee A, Edwards A, Vincentelli R, Cambillau C, Kim R, Kim SH, Rao Z, Shi Y, Terwilliger TC, Kim CY, Hung LW, Waldo GS, Peleg Y, Albeck S, Unger T, Dym O, Prilusky J, Sussman JL, Stevens RC, Lesley SA, Wilson IA, Joachimiak A, Collart F, Dementieva I, Donnelly MI, Eschenfeldt WH, Kim Y, Stols L, Wu R, Zhou M, Burley SK, Emtage JS, Sauder JM, Thompson D, Bain K, Luz J, Gheyi T, Zhang F, Atwell S, Almo SC, Bonanno JB, Fiser A, Swaminathan S, Studier FW, Chance MR, Sali A, Acton TB, Xiao R, Zhao L, Ma LC, Hunt JF, Tong L, Cunningham K, Inouye M, Anderson S, Janjua H, Shastry R, Ho CK, Wang D, Wang H, Jiang M, Montelione GT, Stuart DI, Owens RJ, Daenke S, Schütz A, Heinemann U, Yokoyama S, Büssow K, Gunsalus KC. Protein production and purification. Nat Methods. 2008;5(2):135–46. doi: 10.1038/nmeth.f.202. Erratum in: Nat Methods. 2008;5(4):369. Hallberg, B Martin [added]. PMID: 18235434; PMCID: PMC3178102.

5. Fakruddin M, Mohammad Mazumdar R, Bin Mannan KS, Chowdhury A, Hossain MN. Critical Factors Affecting the Success of Cloning, Expression, and Mass Production of Enzymes by Recombinant E. coli. ISRN Biotechnol. 2012;2013:590587. doi: 10.5402/2013/590587. PMID: 25969776; PMCID: PMC4403561.

6. Rosano GL, Ceccarelli EA. Recombinant protein expression in Escherichia coli: advances and challenges. Front Microbiol. 2014;5:172. doi: 10.3389/fmicb.2014.00172. PMID: 24860555; PMCID: PMC4029002.

7. Rosano GL, Morales ES, Ceccarelli EA. New tools for recombinant protein production in Escherichia coli: A 5-year update. Protein Sci. 2019;28(8):1412–1422. doi: 10.1002/pro.3668. PMID: 31219641; PMCID: PMC6635841.

8. Ki MR, Pack SP. Fusion tags to enhance heterologous protein expression. Appl Microbiol Biotechnol. 2020;104(6):2411–2425. doi: 10.1007/s00253-020-10402-8. PMID: 31993706.

9. Sinha R, Shukla P. Antimicrobial Peptides: Recent Insights on Biotechnological Interventions and Future Perspectives. Protein Pept Lett. 2019;26(2):79–87. doi: 10.2174/0929866525666181026160852. PMID: 30370841; PMCID: PMC6416458.

10. Wang S, Zeng X, Yang Q, Qiao S. Antimicrobial Peptides as Potential Alternatives to Antibiotics in Food Animal Industry. Int J Mol Sci. 2016;17(5):603. doi: 10.3390/ijms17050603. PMID: 27153059; PMCID: PMC4881439.

11. Cui Y, Luo L, Wang X, Lu Y, Yi Y, Shan Y, Liu B, Zhou Y, Lü X. Mining, heterologous expression, purification, antibactericidal mechanism, and application of bacteriocins: A review. Compr Rev Food Sci Food Saf. 2021;20:863–899. doi: 10.1111/1541-4337.12658 PMID: 33443793

12. Sun P, Tropea JE, Waugh DS. Enhancing the solubility of recombinant proteins in Escherichia coli by using hexahistidine-tagged maltose-binding protein as a fusion partner. Methods Mol Biol. 2011;705:259–74. doi: 10.1007/978-1-61737-967-3_16. PMID: 21125392.

13. Waugh DS. The remarkable solubility-enhancing power of Escherichia coli maltose-binding protein. Postepy Biochem. 2016;62(3):377–382. PMID: 28132493.

14. Nguyen AN, Song JA, Nguyen MT, Do BH, Kwon GG, Park SS, Yoo J, Jang J, Jin J, Osborn MJ, Jang YJ, Thi Vu TT, Oh HB, Choe H. Prokaryotic soluble expression and purification of bioactive human fibroblast growth factor 21 using maltose-binding protein. Sci Rep. 2017;7(1):16139. doi: 10.1038/s41598-017-16167-x. PMID: 29170489; PMCID: PMC5700921.

15. Hanahan D. Studies on transformation of Escherichia coli with plasmids. J Mol Biol. 1983;166(4):557–80. doi: 10.1016/s0022-2836(83)80284-8. PMID: 6345791.

16. Kojic M, Strahinic I, Fira D, Jovcic B, Topisirovic L. Plasmid content and bacteriocin production by five strains of Lactococcus lactis isolated from semi-hard homemade cheese. Can J Microbiol. 2006;52(11):1110–20. doi: 10.1139/w06-072. PMID: 17215903.

17. Miljkovic M, Uzelac G, Mirkovic N, Devescovi G, Diep DB, Venturi V, Kojic M. LsbB Bacteriocin Interacts with the Third Transmembrane Domain of the YvjB Receptor. Appl Environ Microbiol. 2016;82(17):5364–74. doi: 10.1128/AEM.01293-16. PMID: 27342562; PMCID: PMC4988209.

18. Miljkovic M, Lozo J, Mirkovic N, O’Connor PM, Malesevic M, Jovcic B, Cotter PD, Kojic M. Functional Characterization of the Lactolisterin BU Gene Cluster of *Lactococcus lactis* subsp. *lactis* BGBU1-4. Front Microbiol. 2018;9:2774. doi: 10.3389/fmicb.2018.02774. PMID: 30498487; PMCID: PMC6249370.

19. Vukotic G, Mirkovic N, Jovcic B, Miljkovic M, Strahinic I, Fira D, Radulovic Z, Kojic M. Proteinase PrtP impairs lactococcin LcnB activity in Lactococcus lactis BGMN1-501: new insights into bacteriocin regulation. Front Microbiol. 2015;6:92. doi: 10.3389/fmicb.2015.00092. PMID: 25713574; PMCID: PMC4322719.

20. Miljkovic M, Malesevic M, Filipic B, Vukotic G, Kojic M. *LraI* from *Lactococcus raffinolactis* BGTRK10-1, an Isoschizomer of *Eco*RI, Exhibits Ion Concentration-Dependent Specific Star Activity. Biomed Res Int. 2018;2018:5657085. doi: 10.1155/2018/5657085. PMID: 29789800; PMCID: PMC5896346.

21. Malesevic M, Stanisavljevic N, Miljkovic M, Jovcic B, Filipic B, Studholme DJ, Kojic M. The large plasmidome of Lactococcus lactis subsp. lactis bv. diacetylactis S50 confers its biotechnological properties. Int J Food Microbiol. 2021;337:108935. doi: 10.1016/j.ijfoodmicro.2020.108935. PMID: 33152568.

22. Raran-Kurussi S, Waugh DS. Expression and Purification of Recombinant Proteins in Escherichia coli with a His_6_ or Dual His_6_-MBP Tag. Methods Mol Biol. 2017;1607:1–15. doi: 10.1007/978-1-4939-7000-1_1. PMID: 28573567; PMCID: PMC7122414.

23. Bucher D, Grant BJ, McCammon JA. Induced fit or conformational selection? The role of the semi-closed state in the maltose binding protein. Biochemistry. 2011;50(48):10530–9. doi: 10.1021/bi201481a. PMID: 22050600; PMCID: PMC3226325.

